# Reproducing Human Motor Adaptation in Spiking Neural Simulation and known Synaptic Learning Rules

**DOI:** 10.1101/2021.06.24.449760

**Authors:** Yufei Wu, Shlomi Haar, Aldo Faisal

**Affiliations:** Brain and Behaviour Lab, Dept. of Bioengineering and Dept. of Computing, Imperial College London, London, UK; Department of Brain Sciences and UK Dementia Research Institute – Care Research & Technology, Imperial College London, London, UK; Behaviour Analytics Lab, Data Science Institute, Imperial College London, London, UK; UKRI Centre for Doctoral Training in AI for Healthcare, Imperial College London, London, UK; MRC London Institute of Medical Sciences, Imperial College London, London, UK

## Abstract

Sensorimotor adaptation enables us to adjust our goal-oriented movements in response to external perturbations. These phenomena have been studied experimentally and computationally at the level of human and animals reaching movements, and have clear links to the cerebellum as evidenced by cerebellar lesions and neurodegeneration. Yet, despite our macroscopic understanding of the high-level computational mechanisms it is unclear how these are mapped and are implemented in the neural substrates of the cerebellum at a cellular-computational level. We present here a novel spiking neural circuit model of the sensorimotor system including a cerebellum which control physiological muscle models to reproduce behaviour experiments. Our cerebellar model is composed of spiking neuron populations reflecting cells in the cerebellar cortex and deep cerebellar nuclei, which generate motor correction to change behaviour in response to perturbations. The model proposes two learning mechanisms for adaptation: predictive learning and memory formation, which are implemented with synaptic updating rules. Our model is tested in a force-field sensorimotor adaptation task and successfully reproduce several phenomena arising from human adaptation, including well-known learning curves, aftereffects, savings and other multi-rate learning effects. This reveals the capability of our model to learn from perturbations and generate motor corrections while providing a bottom-up view for the neural basis of adaptation. Thus, it also shows the potential to predict how patients with specific types of cerebellar damage will perform in behavioural experiments. We explore this by *in silico* experiments where we selectively incapacitate selected cerebellar circuits of the model which generate and reproduce defined motor learning deficits.

**Author summary:** A rich body of work in human motor neuroscience developed high-level computational theories of sensorimotor control, learning and adaptation. But there is a gap in understanding how this may be implemented and learn on the level of neurons, synapses and spikes. Conversely, studies of patients with cerebellar lesions or neurological disease highlight the essential role the cerebellum plays in our ability to perform motor learning. Yet, how these anatomical and molecular defects play out in terms of human movement have to date not been linked to a model that spans multiple level of biological organisation from neural circuits to reproducing human motor experiments. To address this gap, we present a spiking neuron of the sensorimotor system focused on the cerebellum, with which we can on the one side reproduce the high-level behaviour learning phenomena observed in healthy subjects, as well as quantitatively predicting the putative effects on human movement trajectories of cerebellar lesions implemented at the cellular level.

## Introduction

Human behaviour is sometimes influenced by external perturbations in the environment, which cause unexpected deviation from the desired movement and may dramatically influence movement accuracy and stability. Due to the large delay in sensory feedback, it is very hard to respond immediately to the deviation and correct the current movement within the execution period. Nevertheless, the brain shows its capacity of generating efficient compensations against the external perturbation after repeated exposure to the same environmental conditions. This is known as adaptation or error-based learning [1], a key element of sensorimotor learning in the brain. It enables accurate execution of movements under mechanical perturbations [2,3] via error-based movement modifications.

Sensorimotor adaptation is strongly hypothesised to have neural correlates of the cerebellum [4–10]. Traditionally, the cerebellum is considered to contain an error-driven learning system (Marr-Albus theory [11,12]) and to monitor motor commands according to internal feedback [13], using a motor-to-sensory predictive model (the forward model [14–16]). This internal feedback (sensory predictions) accurately follow the actual body state if the forward model is well trained. Sudden exposure to unexpected perturbations leads to sensory prediction errors, which update the forward model for new dynamics. From a systematic perspective, this forward model forms a predictive control [17] system in the cerebellum, modifying the motor commands in order to control the complex system of the body with high-order dynamics and large sensory feedback delays.

Sensorimotor adaptation is typically studied in basic sensory-driven tasks with repeated trials, e.g. reaching [18], throwing [19] and walking [20]. Each adaptation procedure is summarised by a trial-by-trial curve revealing the learning feature of error-based learning, which is normally explained using state-space models. A simple single-state model is capable of capturing exponential-like curves during adaptation and wash-out, but a two-state model [21], involving two independent learning processes (states) with different learning and decaying rate, is a more convincing model successfully explaining important sensorimotor adaptation features like savings [22–24] and aftereffect [18,19]; and it provides a good fit even for motor learning in the real world [25]. Nevertheless, state-space models show limitations of being top-down approaches: 1) trial-level extractions of the adaptation procedure ignores the time-continuous learning effects and body-environment dynamics within trials; 2) the mapping from different learning processes (states) [26] onto neural substrate is unclear, as anatomical and cellular mechanisms do not constrain these abstract models; 3) it lacks theoretical support to choose the appropriate number of states while keeping a balance between the fitting performance and model complexity. Due to these limitations, state-space models are more suitable for studying the behaviour (consequences of motor control) instead of learning mechanisms in the brain.

Bottom-up models can be useful to better understand the internal mechanisms of sensorimotor adaptation in the brain. Here, we present a spiking neural network with synaptic learning rules to simulate error-based learning from the neuronal level to the behavioural level. This model follows the biological structure (Marr-Albus theory) and functional structure (the forward model) of the sensorimotor adaptation with populations of spiking neurons representing different components of the cerebellum. The predictive learning in the forward model is achieved via the co-action of long-term depression (LTD) and long-term potentiating (LTP) synaptic learning of Purkinje cells [27] under predictive error signals. On this bases, we add another learning mechanism of memory formation [28,29] with simple Hebbian synaptic learning, which can be considered as a memory-based extension of the forward model. Unlike state-space models, our model provides simultaneous monitoring of neurons and movement trajectories throughout each trial.

## Materials and methods

### Mechanism and Signal Flow of Sensorimotor Adaptation

Let 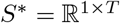 be a planned voluntary movement represented by a time-series of body states. 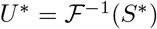 is the corresponding motor command in the brain based on the target movement, where 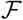 represents the kinematics and dynamics of the body. If both motor planning and execution are accurate, the executed motor command *U* = *U*\* and the observed body state *S* = *f*(*U*) = *S*\* matches the expectation perfectly.

When an unexpected perturbation *P* is introduced to the system causing movement deviation *g*(*P*), the observed body state then reflects the combining consequence of motor execution and perturbation 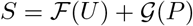. Motor adaptation is a procedure of modifying executed motor command *U* for compensating 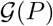. We assume that this modification is achieved by adding an additional compensation signal *U*′ (*U* = *U*\* + *U*′) instead of updating *U*\*. Fig. 1 explains the mechanism of the cerebellum-based adaptation, which is later applied in our spiking neural network simulation. Let [*n, t*] represents the t-th time point in n-th trial. We describe the adaptation procedure as generation of motor compensation *U*′[*n, t*] under two learning processes - forward model and memory formation.

**Fig 1.**
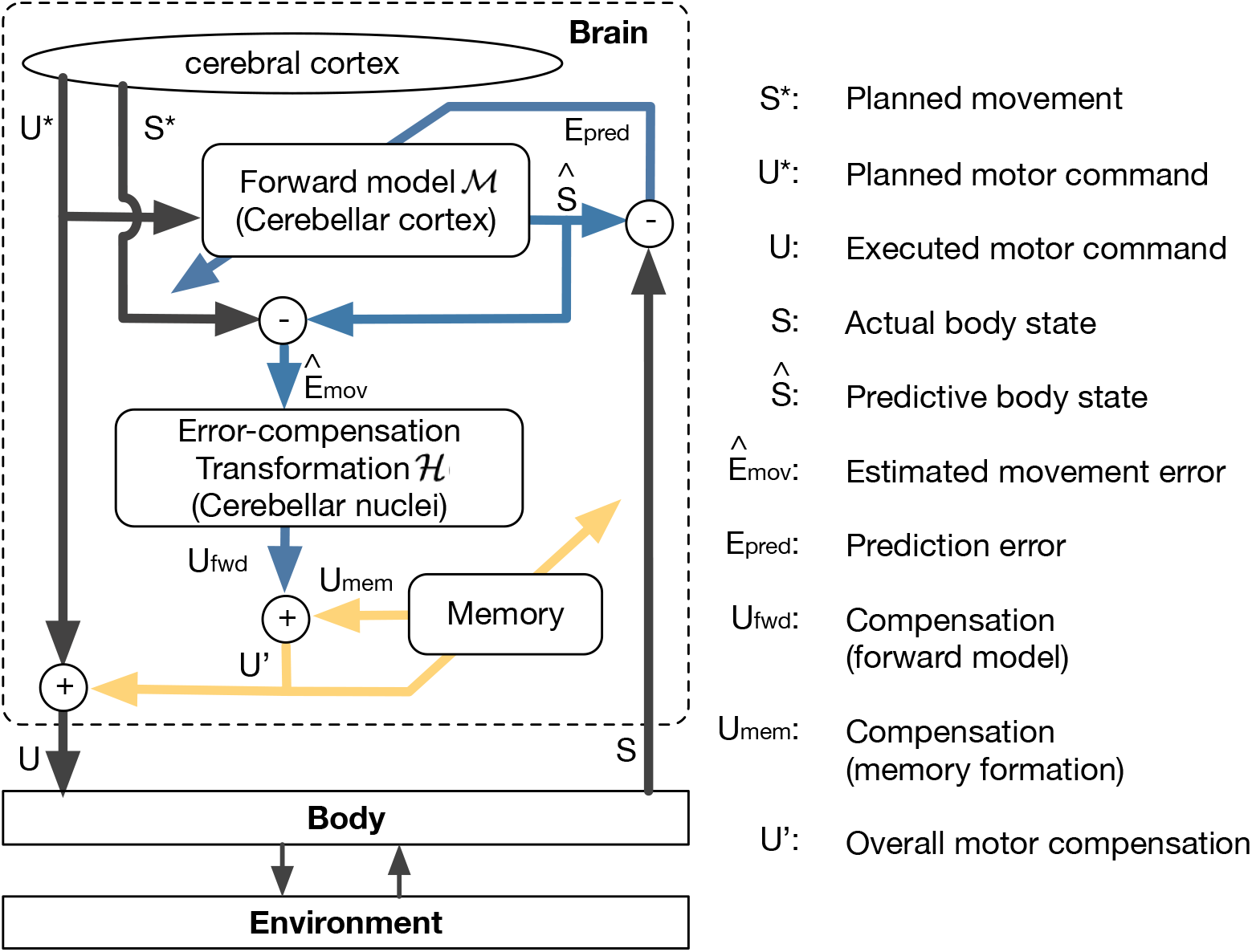
Graphical structure of cerebellum-based adaptation mechanism. Motor compensation *U*′ comes from the co-action of predictive learning in the forward model (blue arrows) and the memory formation (yellow arrows).

Let 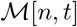 represents the forward model at a particular time step. It receives efference copy *U*\*[*t*] and provides the predictive body state 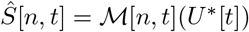. This predictive body state is involved in the calculation of two different error signals (see blue arrows in Fig. 1):

- The estimated movement error 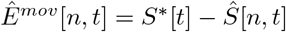, which triggers the compensation signal *U_fwd_*[*n, t*] with transformation 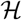:

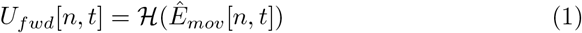
- The prediction error 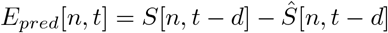 considering the sensory feedback delay *d*. This is the teaching signal for updating the forward model:

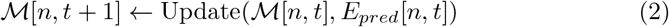 The memory formation relies on the compensation signal generated by the forward model. As a memory process, it gradually consolidates previous motor compensation signals (see yellow arrows in Fig. 1) in the same stage of the movement:

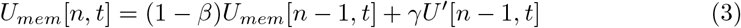

where *β* refers to the decay rate, which is usually small to show resistance to forgetting and *γ* is the speed of transformation to the long-term memory. This memory process provides a empirical support *U_mem_*[*n, t*] for the generation of overall motor compensation *U*′[*n, t*]:

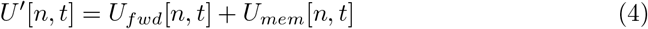

### Cerebellum-based Spiking Neural Network for Sensorimotor Adaptation

Following the mechanism explained in Fig. 1, we present a spiking neural network implementation of sensorimotor adaptation. This spiking neural network (Fig. 2 (A)) contains three main components: input neurons, the cerebellar cortex and the cerebellar nuclei. Input neurons encode upstream signals (planned movement *S*\* and planned motor command *U*\*) and spiking activities of input neurons are carried by mossy fibres, projecting to the cerebellar cortex and the cerebellar nuclei (green arrows).

**Fig 2.**
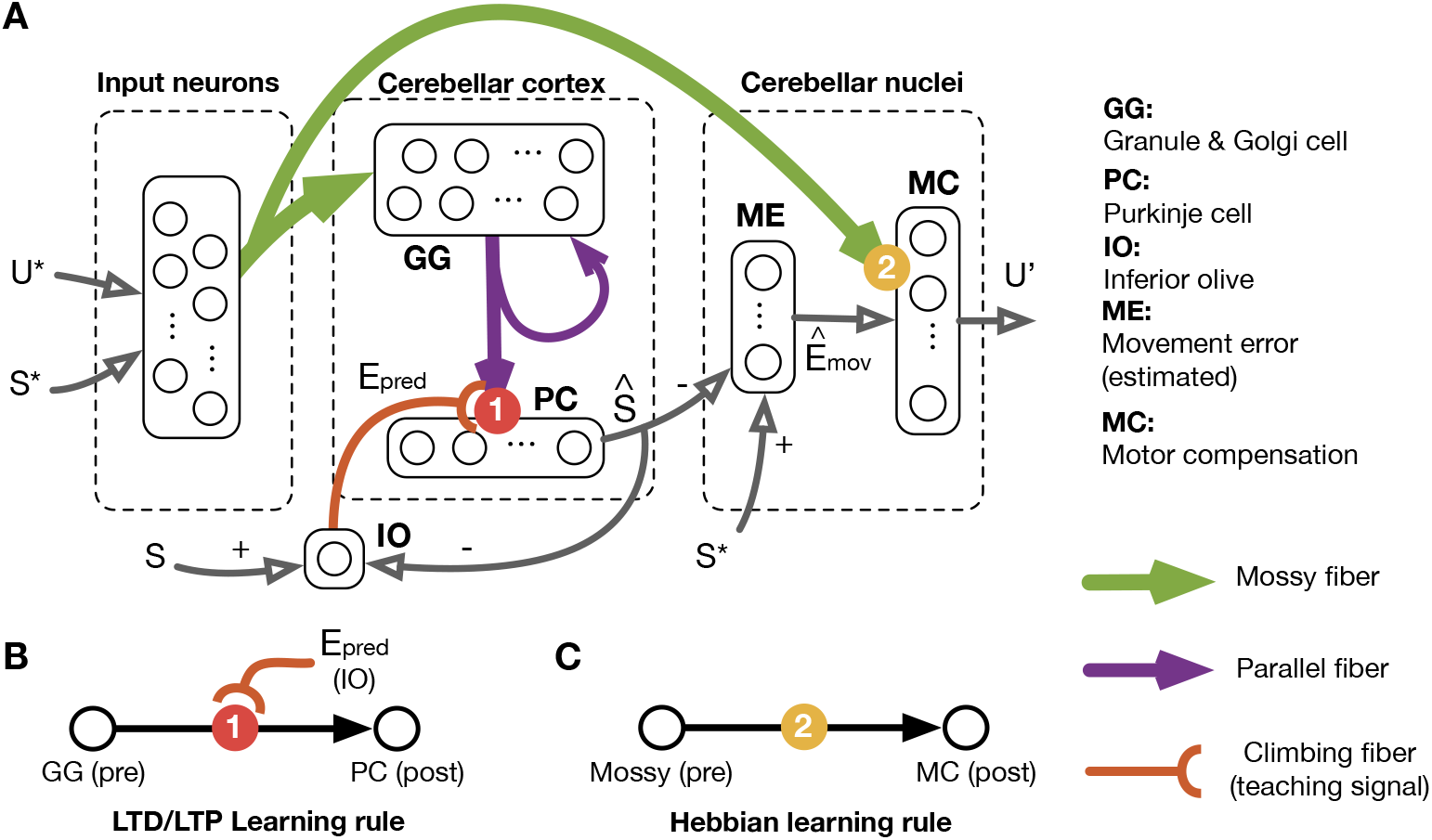
(**A**) Cerebellum-based spiking neural network for sensorimotor adaptation, consisting of input neurons, the cerebellar cortex and the cerebellar nuclei. Each component involves populations of spiking neurons linking with synapses. Circles with numbers represent two synaptic learning rules applied in this spiking neural network for adaptation. **B**) Spiking neural model and population size of neural populations in the cerebellum-based spiking neural network. Poisson: Poisson neurons. Izhikevich RS: Izhikevich regular spiking neurons [30]. IAF: integrated-and-fire neurons. (**C**) LTD/LTP synaptic learning rules with prediction error *E_pred_* as the teaching signal, which takes place between parallel fibres and Purkinje cells (see **A**). (**D**) Hebbian synaptic learning rule for memory formation, which takes place between mossy fibres and the motor compensation neurons in cerebellar nuclei (see (**A**).

The cerebellar cortex in our model is a spiking neural implementation of the forward model, which contains two populations of spiking neurons representing the granule cell layer and the Purkinje cell layer. *In vivo*, the granule cell layer involves mainly excitatory granule cells and inhibitory Golgi cells, receiving signals from mossy fibres and projecting to the Purkinje cell layer via parallel fibres. Thus, we introduce random inhibitory recurrent connections to the granule/Golgi population, which provides complex spatial-temporal firing patterns depending on all previous inputs and the correlation between firing patterns decay over time [31]. These firing patterns innervate Purkinje cells via parallel fibres (purple arrows), providing estimations of the current body state 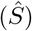. These two spiking neural populations form a liquid-state machine with the granule layer as the recurrent network and the Purkinje cell layer as the read-out neurons.

The cerebellar nuclei in our model are designed to involve two populations of spiking neurons. The first population encodes the estimated movement error 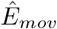 as the difference between the Purkinje read-out (estimated body state 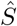) and the planned body-state as part of the movement *S*\*. The other population, referred as motor compensation, is innervated by signals from two pathways: 1) the estimated movement error signal from the first population inside the cerebellar nuclei; 2) the integrated signal from mossy fibres. The former can be considered as the error-compensation transformation 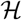, while the latter is the compensation memory *U_mem_*. The combination of input stimuli from these two pathways produces the final motor compensation output *U*′ of the motor compensation population. Fig. 2 (B) illustrates the spiking neural model, and the size ratio between the different neural populations. More implementation details can be found in *Appendix*.

During the simulation, we need to update the strength of synaptic connectivity to enable predictive learning and memory formation in this cerebellum-based spiking neural network. We apply two different synaptic learning rules (see Fig. 2 (A-C)):

#### Error-based LTD/LTP Synaptic Learning Rule for Predictive Learning

The error-based LTD/LTP learning rule is applied to update synaptic weights between parallel fibres and Purkinje cells. At a certain time point, these synapses experience either LTP or LTD depending on the stimuli of the climbing fibre [32], carrying the prediction error signal *E_pred_*. Large errors lead to LTD so that synaptic connections causing deviated predictions are weakened. Otherwise, these synapses would remain updating with LTP, which strengthens corresponding synaptic connections. This LTD/LTP synaptic learning is modelled by a 3-factor updating rule (see Fig. 2 (B)), depending on 1) pre-synaptic (parallel fibres) firing activities, 2) post-synaptic (the Purkinje cell layer) firing activities and 3) error-based teaching signal of predictive error. The following equations demonstrate the details of the weight update rule:

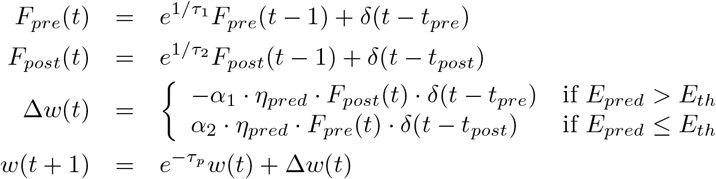

where *w* is the synaptic weight, *t_pre_* and *t_post_* refer to the spiking times of the pre-synaptic neuron and the post-synaptic neuron respectively. *η_pred_* is the learning rate and *δ*(*x*) equals to 1 if *x* = 0 and 0 otherwise. *τ_p_, τ*_1_, *τ*_2_ are time constants and *α*_1_, *α*_2_ are positive factors determined by the predictive error *E_pred_*.

#### Hebbian learning for memory formation

Previous experience of adaptation is consolidated into memory via the Hebbian synaptic learning rule (see Fig. 2 (C)) applied to synapses between mossy fibres and motor compensation neurons. In our model, these synapses correspond to the memory system in Fig. 1, transferring motor compensation into long-term memory in the cerebellar nuclei. This synaptic updating rule is described as:

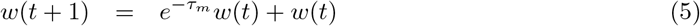

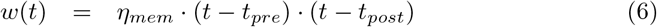

where *w* is the synapses between a pre-synaptic input neuron and a post-synaptic neuron in the cerebellar nuclei network, whose spiking activities are marked as *t_pre_* and *t_post_* respectively. *η_mem_* is the learning rate and *τ_m_* is a decay constant, showing that memory is gradually decaying.

## Results

We tested the cerebellum-based spiking neural network in the force-field paradigm [2], which is a reaching task in a planar surface (x-y plane) under external force perturbations. The target point is placed 8 cm ahead from the starting point (Fig. 3 (A)), and the intention is to make straight hand movements reaching the target point. We introduced four different settings of force-field trials, common in force-field adaptation studies (Fig. 3 (B)): 1. baseline trials (BT) with no external perturbations; 2. adaptation trials (AT) with external forces, parallel to the x-axis towards the right side and proportional to y-axis velocities of hand movements; 3. de-adaptation trials (DT) with same perturbation settings as the adaptation task but towards the left-hand side; 4. error-clamp trials with all x-axis deviation clamped by external forces. With these four types of trials, we designed various motor adaptation experiments protocols, previously used in human studies and state-space models [21]. Before each experiment, our spiking neural network was pre-trained in baseline trials until it can generate stable predictions with the forward model and execute vertical movement under baseline conditions.

**Fig 3.**
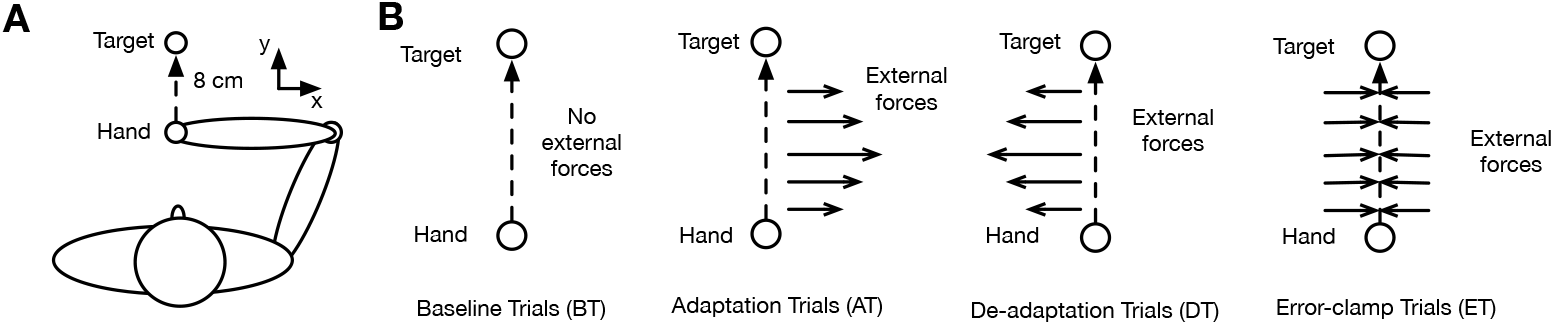
Experimental design of force-field trials. (**A**) Task description. The aim of the task is to move the hand forward from a start point to a target point, which is 10 cm ahead of the start point. (**B**) Different types of force-field trials. External forces in adaptation and de-adaptation trials are proportional to the y-axis velocity.

To better evaluate the effect of adaptation, we apply an adaptation coefficient *K*1 as a metric to measure the force-field adaptation performance [21], which is a regression coefficient representing the sensitivity of the actual human-produced force pattern proportional to the ideal compensatory force pattern.

### Movement Correction and Aftereffect

We tested the adaptation capability of our spiking neural network in an experiment containing an adaptation phase of 70 adaptation trials (AT) followed by a wash-out phase of 20 baseline (BT) trials (Fig. 4 (A)). Fig. 4(B) present the adaptation coefficient of each trial during this experiment (averaged over 10 simulations). When our model first encounters the unexpected external perturbation, it responses quickly to cancel the influence of the perturbation by generating effective compensatory forces. Fig. 4(C) shows a window of this ‘competition’ between the external perturbation and internal motor compensation from spiking activities to the movement trajectory. As a reflection, the adaptation coefficient curve shoots up in the first several trials. As the spiking neural network experiences more adaptation trials, the adaptation coefficient continues to incline, at a relatively lower rate, until it reaches a stable status, showing an exponential-like learning curve during the adaptation stage of the experiment. During the wash-out stage (baseline trials after adaptation), the external perturbation suddenly vanishes. The adaptation coefficient remains high at the very beginning and then gradually decreases to around zero as the network is exposed in more wash-out trials. This phenomenon is regarded as the aftereffect (see arrow in Fig. 4(B)), which is also observed in human sensorimotor adaptation experiment (e.g. [33]).

**Fig 4.**
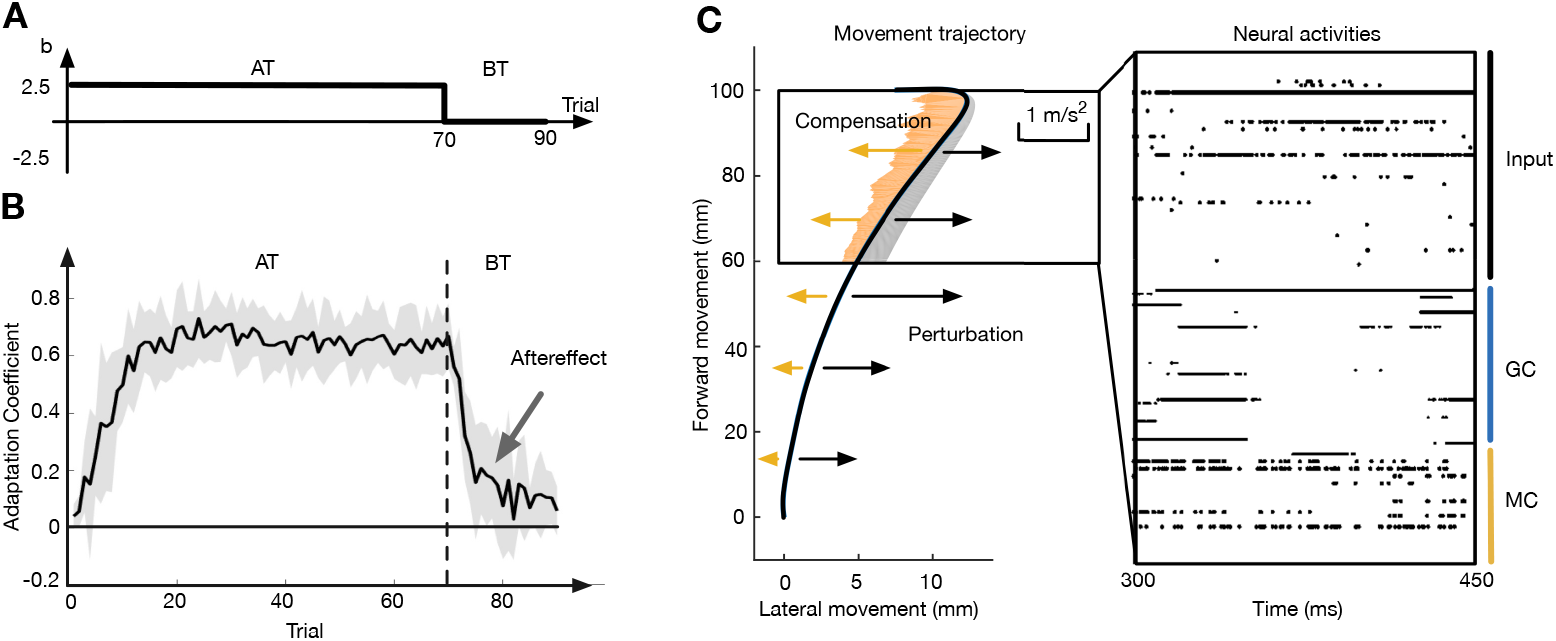
(**A**) Experimental design of sensorimotor adaptation experiments involving 70 adaptation trials (AT) and 20 baseline trials (BT). b is the ratio between the force perturbation and the y-axis velocity. (**B**) Adaptation coefficient during the adaptation experiment in (A), averaged over 10 simulations. (**C**) A window showing the external perturbation (grey shaded bar), the motor compensation (yellow shaded bar) and corresponding neural activities during a single adaptation trial. Input: input neurons. GC: (part of) Granule cells. MC: motor compensation neurons.

Taking advantages of spiking neural simulations, one can observe detailed neural activities and movement trajectories on a trial-by-trial basis at the different stages of an adaptation experiment. We choose 5 sample trials from one specific simulation (Fig. 5(B)) to represent different adaptation stages and present movement trajectories during these trials in Fig. 5(B). Fig. 5(C) present the neuronal activities and their decoded information during these trials which induced the observed behaviour. In trial 1, the external perturbation is completely unexpected. The estimated hand state ([*x, ẋ*) remains the same as baseline trials and delivers zero estimated movement error to the compensation generator, which generates no compensation. The movement trajectory deviates from the straight path towards the target and terminates on the right-hand side with a large error (48mm). In this condition, the prediction error is large and triggers the update of the forward model until its estimation matches the actual body state. As the forward model in the spiking neural network provides a reasonable estimation of the movement error, motor compensation signals are generated correspondingly. During the next several trials, our model responses quickly and effectively against the perturbation in adaptation trials, and as a result, the end-point error decreases to !20 mm in Trial 12.

**Fig 5.**
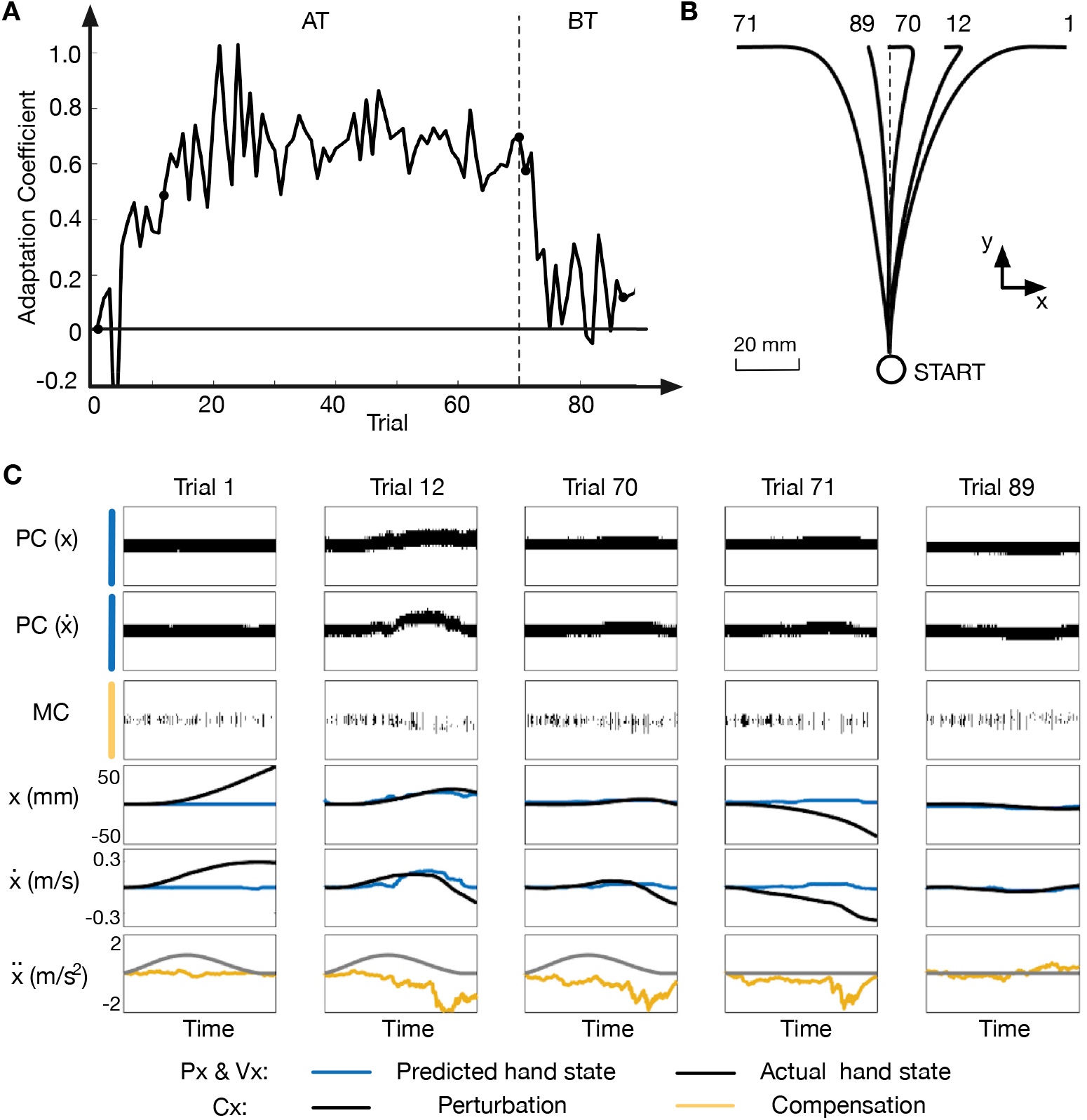
Neural activities and movement trajectory during an adaptation experiment. (**A**) Adaptation Coefficient in an experiment with 70 adaptation trials and 20 baseline wash-out trials. (**B**) Movement trajectories in different trials. (**C**) Selected raster plots of neural activities during 5 different trials: two groups of Purkinje cells (PC) representing x-axis position (*x*) and velocity (*ẋ*) and one group of motor compensation neurons (MC). Each raster plot represent neurons across a neural population (across the vertical axis), Each row of dots correponds to one neuron and we have organised the order of neurons along the vertical axis based on their receptive field placement. Therefore e.g. movements induce adjacent neurons in the raster plot to become active and inactive which is visible by the broad black brand of activity involving many neurons following e.g. the black band in the PC plots which tracks the blue curve in the lower plots.

The observed amplitude of motor compensation is much larger in the second half of the movement. The reason is that the movement error is not obvious at the beginning of the movement since the effect of perturbation is not yet accumulated. As the adaptation continues, our model achieves a stable stage eventually - the predictive model follows the actual trajectory, leading to subtle movement error. Although the forward model contributes little to the motor compensation at this stable stage, the adaptation preserves (trial 70 in Fig. 5(C)) due to the existence of memory formation. In trial 71, the disappearance of external forces lead to the aftereffect and the remaining motor compensation drives the hand in the opposite direction. The aftereffect is washed away quickly and the spiking neural network returns back to its initial state with a straight trajectory generated towards the target point (5 (B) Trial 89).

### Memory Effect in Adaptation

In this part, we present two more experiments for studying the memory effect of our spiking neural network during sensorimotor adaptation, namely the learning-relearning experiment and the error-clamp experiment.

The learning-relearning experiment involves two phases of adaptation trials. The first phase of 70 adaptation trials (learning phase) matches the experiment in Fig. 4 and the second adaptation phase of 55 adaptation trials (relearning phase) is a re-exposure to the same condition after a short interval of 5 de-activation trials (see Fig. 6 (A)). Fig. 6(B) shows the averaged adaptation coefficient (N=10) during this learning-relearning experiment. During de-adaptation trials, the adaptation coefficient decreases rapidly. When the relearning starts after the end of de-activation trials, the adaptation coefficient eventually reaches below baseline, meaning that our model has already acquired some learning experience in an opposite condition. Nevertheless, our model shows a much faster adaptation ability in relearning as the adaptation coefficient immediately jumps back to a high level. This sudden jump is caused by the preserved memory of motor compensated at the beginning of the relearning phase and the same phenomenon in human adaptation is referred to as ‘savings’ [34,35].

**Fig 6.**
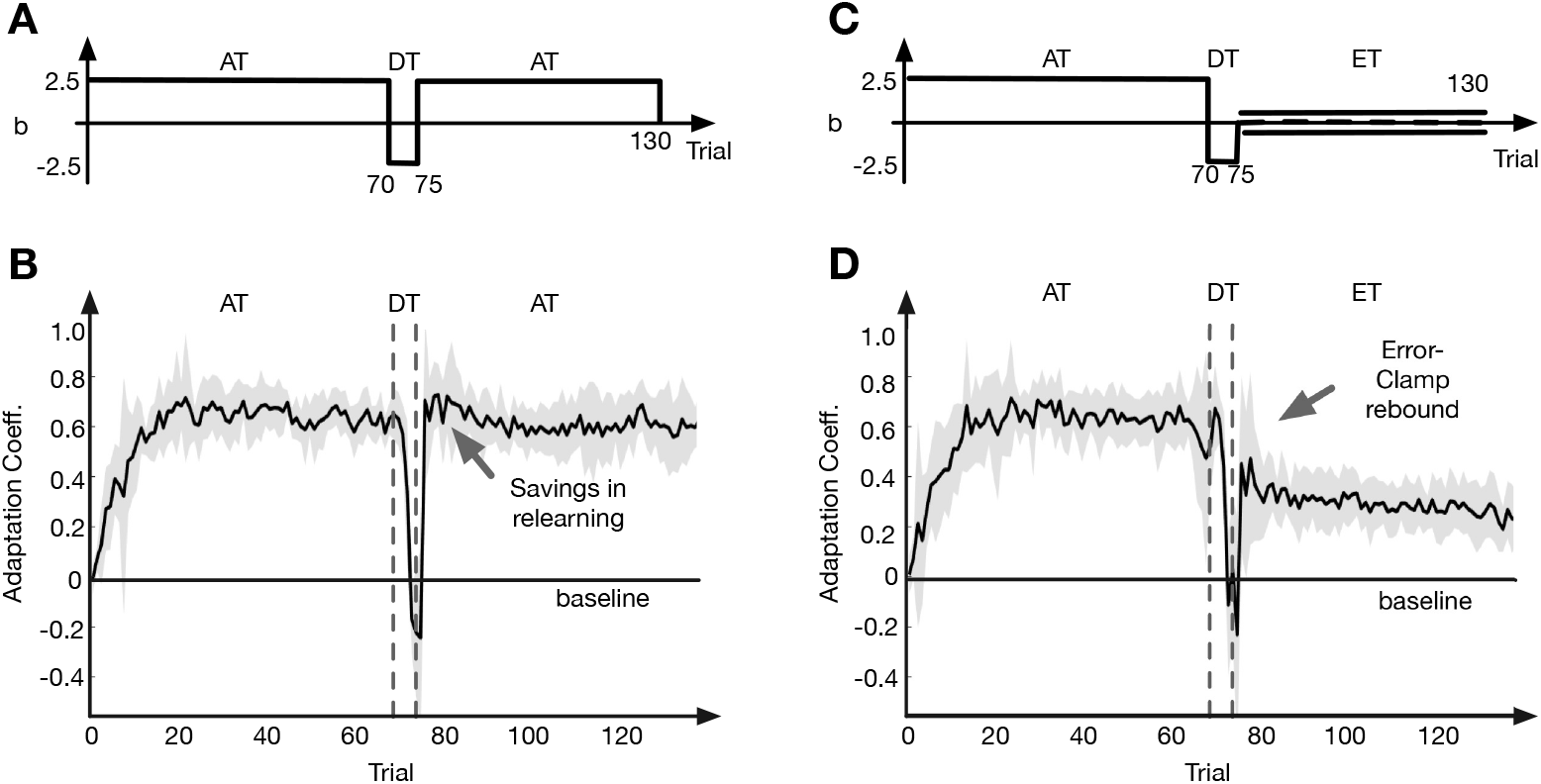
Memory effect in adaptation. (**A**) Experimental design of learning-relearning experiment, involving 70 adaptation trials (AT), 5 de-adaptation trials (DT) and another 55 adaptation trials for relearning. (**B**) Adaptation performance of the cerebellum-based spiking neural network in the learning-relearning experiment using our cerebellar learning spiking model (N=10). Savings are observed in the relearning phase. (**C**) Experimental design of the error-clamp experiment, involving 70 adaptation trials (AT), 5 de-adaptation trials (DT) and 55 error-clamp trials (ET). (**D**) Adaptation performance of the cerebellum-based spiking neural network in the error-clamp experiment (N=10). The adaptation coefficient rebounds above baseline in error-clamp trials.

In order to further reveal the adaptation retention caused by memory formation, we introduce the error-clamp experiment by replacing the relearning phase with an error-clamp phase containing 55 error-clamp trials (Fig. 6(C)). In error-clamp trials, movements are restricted to follow a straight-forward routine towards the target. However, we can still estimate the equivalent ‘lateral deviation’ from applied lateral forces as if the restriction does not exist and thus calculate the adaptation coefficient. We can observe a rebound phenomenon of the adaptation coefficient at the beginning of the error-clamp phase (see arrow in Fig. 6(D)). Since the movement error in error-clamp trials is always zeros, motor compensation generated by the predictive learning quickly vanishes and the rebound of the adaptation coefficient mainly reflect the effect of remaining memory of motor compensation, which is also captured in human experiments (e.g. [23]). After this rebound, the adaptation coefficient decays slowly over time showing its resistance to forgetting.

### From Cerebellar Lesion to Behaviour Impairment

Damage to the cerebellum may cause severe dysfunctions in motor control of smooth and accurate movements and motor adaptation to novel dynamics [36]. Patients with cerebellar disorders show various levels of performance decease in sensorimotor adaptation tasks, from less steep learning curves to little reservation of adaptation ability [37–42]. Our model enables asking questions about how different cerebellar lesions or neurodegeneration might affect adaptation. With our bottom-up spiking neural simulation of the cerebellum, we can provide predictions to answer these questions by manually introducing lesions to the network.

We introduced four different lesions to the cerebellar circuitry of our cerebellum-based spiking neural network. We tested these lesioned networks in the same experiment shown in Fig. 4 (A), consisting of 70 adaptation trials and 20 baseline trials, and present the adaptation performance with comparison to a control network with no lesion introduced (Fig. 7. The first lesion models neurodegeneration in normal aging, which leads to atrophy of about 20% of the cerebellar gray matter by the age of 80 [43,44]. Accordingly, we lesioned 20% of the neurons in all layers of our model. As a result of this lesion we see a decay in the adaptation coefficient which plateau earlier and an increase in the inter-subject variability. This is in line with the effect of aging reported in force-field adaptation [45] as well as in visuomotor rotation adaptation [46] and in split-belt walking adaptation [47]. Such effect was also reported in few cerebellar patients studies [10, 40, 42]. The second lesion was to the mossy fiber terminals in the DCN, which are known to strengthen with learning, thus related to motor memory formation in the cerebellum [29, 48]. Here, we lesioned all the mossy fiber projections to the DCN in our model. The lesion led to a decay in the adaptation coefficient and plateau – similar to the one reported in the previous model – but without any increase in the inter-subject variability. Notably, in this lesion model the washout is faster than in the control while in the previous one, which actually makes this model more in line with older-adults [45] and cerebellar patients [42] performance. The third lesion was a selective loss of Purkinje cells, previously reported in cases of cerebellar ataxia [49, 50]. Accordingly, we lesioned 50% of the Purkinje cells in our model. This PC only lesion led to slowing down of learning, but learning continues over the entire adaptation phase towards the learning plateau of the non-lesioned model. Such learning curve was indeed reported in few cerebellar patients studies [37,40]. The fourth “lesion” was an increased noise level of Purkinje cell. Effectively neurons are finely balanced in their mechanisms so as to reduce the impact of their internal and external noise sources on their signalling. Any intermediate imbalance due to disease or damage, will thus likely result in noise increases, which will effect neural function [51]. There are multiple ways by which neuronal and synaptic noise may dramatically increase pathologically [52], such as neurodegeneration or inflammation [53, 54] or mutations in channelopathies [55]. In mice models, increase in noise as result of an voltage-gated calcium channels imbalance was shown to induce ataxic behaviour [56]. Given that there are therefore multiple mechanisms that may increase noise and its relation to ataxic behaviour, a “noise lesion” was applied in the model by dramatically increasing the background white noise of Purkinje cells (10 times the original amplitude). This lesion, like the first, induced a decay in learning which lead to a lower adaptation plateau, an increase in the inter-subject variability and a decay in the washout as well. Here the increase in the inter-subject variability was much more moderate than in the first, while the decay in the adaptation and washout was much more severe – almost flattening the learning curve. Such learning curve resembles the lack of adaptation and washout reported for some cerebellar degeneration patients [39].

**Fig 7.**
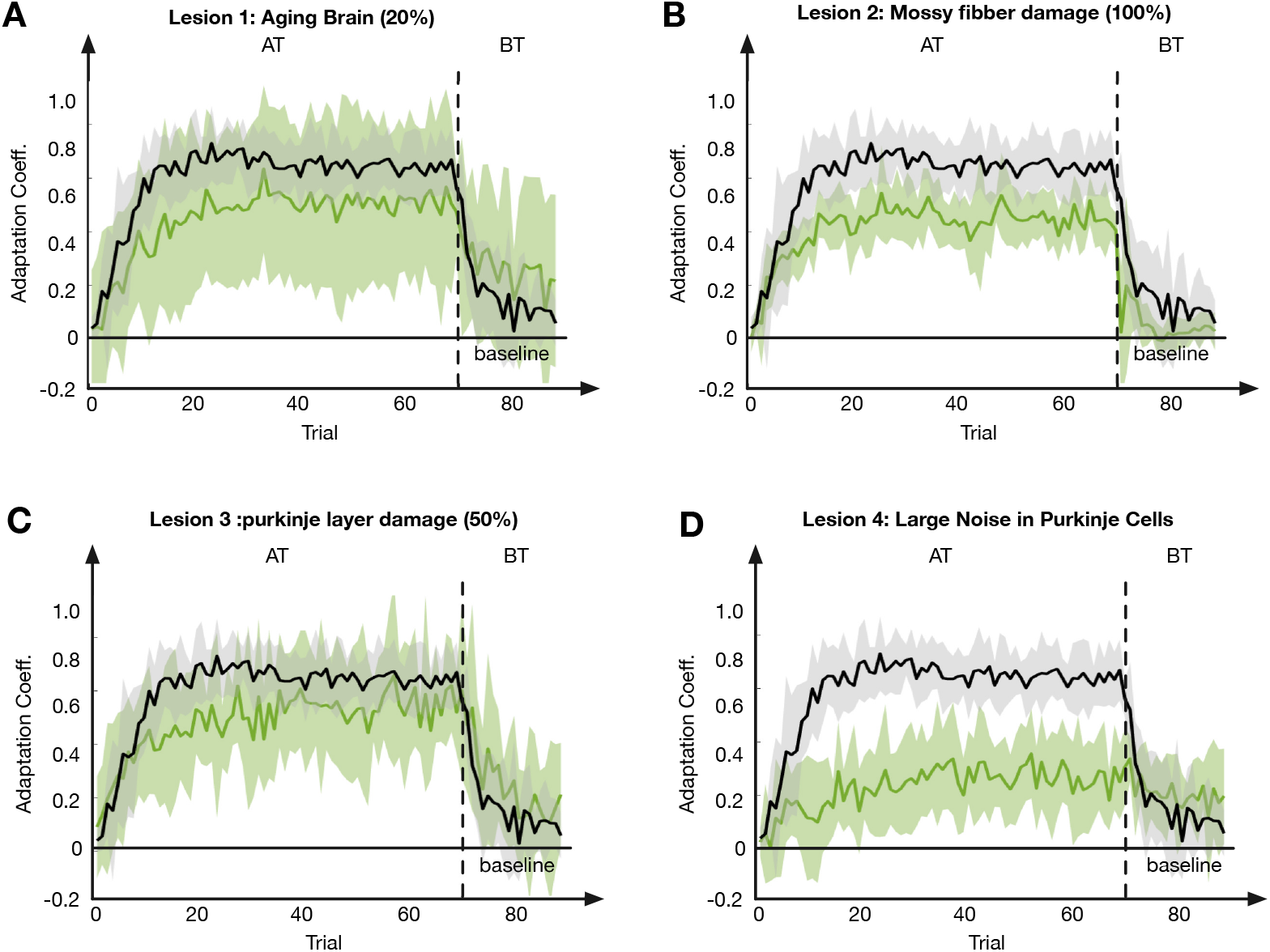
Adaptation performance under 4 kinds of cerebellar lesions (N=10) in an experiment with 70 adaptation trials (AT) and 20 baseline trials (20). Black curve: control. Green curve: (**A**) Lesion 1: Neurodegeneration in aging brain. (**B**) Lesion 2: Mossy fiber damage. (**C**) Lesion 3: Purkinje layer damage. (**D**) “Lesion” 4: large noise increase in Purkinje layer.

## Discussion

In this paper, we built a spiking neural network modelling the human sensorimotor adaptation. This spiking neural network spans several orders of magnitude of biological organisation, from individual cerebellar neurons and synapses to the trajectory of hand reaches under various force perturbations. To test the capability of our model in sensorimotor adaptation tasks, we designed a series of experiments containing different types of force-field trials. The results demonstrated that our model’s ability to learn from errors and generate motor compensations to correct movements against external perturbation. In addition, the adaptation performance of our model showed various learning phenomena observed in human adaptation tasks, e.g. aftereffects, savings in relearning and error-clamp rebounds. By introducing different types of “lesions” to the spiking neural networks model, we present how cerebellar lesions may damage the ability of sensorimotor adaptation from the bottom-up perspective.

Conceptually, our cerebellum-based spiking neural network is similar to the multi-rate learning model [21], which explains interactions systematically with multiple components [21, 26, 57–61]. In our spiking neural network, the predictive learning learns fast but shows no ability of retention, which matches the description of the ‘fast learning’ process in the multi-rate learning model. By contract, the compensation memory, corresponding to the ‘slow learning’ process is formed slowly but is resistant to decaying and washing-out. From this perspective, our model also provides extensions and improvements to abstract state-space models with 1) biological similarities; 2) synaptic learning rules; 3) time-continuous simulations within trials.

Many phenomena in sensorimotor adaptation are consequences of memory, e.g. savings and error-clamp rebounds. It is suggested that memory formation is involved in sensorimotor adaptation (e.g. [28, 62]), the brain is capable of remembering and recalling things learned previously. However, it remains unclear where and how the task-based memory is formed in the brain. In computational models, the memory mechanism is usually explained via low decay rates of learning processes [21,61] or memory of errors formed as the belief of the new environment [63]. The memory formation in our model, which is different from those mentioned above, remembers directly from previously generated compensations. This is due to the error-compensation conversion which is similar to a black-box in force-field tasks - one can not directly estimate the ideal force compensation from the observed error. Our solution of compensation memory formation tackles the problem without the additional requirement of an inverse model.

### Gradient Descent vs Synaptic Plasticity

Two main aspects that set our framework are: 1) no dependency on gradient calculations or back-propagation; 2) functioning on partially observed models with hidden black-box components. The first aspect leads to an open question on ‘if the brain implements error back-propagation and how’. Directly applying gradient-based modifications for synaptic connections are biologically unrealistic [64]: neurons do not have double-directional reciprocal connections carrying forward stimuli and backward gradients. Recent models, e.g. temporal-error models and explicit-error models, provide approximate solutions for back-propagation in with synaptic rules [65–69]. However, these models require explicit patterns of the desired output in the neuronal level, which is not feasible and brings to our second statement of black-box components. The entire system determining body movements involves black-box components like body kinematics and the environment while no information can be back-propagated through those black-boxes to provide gradients or stimulate desired patterns.

### Cerebellar Lesions and simulation

Motor adaptation studies in animal models can validate the prediction of our model by creating similar lesions in the animal model and the computational model and compare the resulting behaviours. For example, a motor adaptation study in zebrafish showed decay in motor adaptation following lesions to the inferior olive [70]. This is overlapping with lesion 1 described here. At the same time, comparing the predictions from the model lesions to the behavioural results of cerebellar patients can shed new light on the neural mechanisms behind their symptoms. To date, motor adaptation studies mostly described the patients as one group, averaging across them (e.g. [37–39, 42]), while only few split the patients to groups based on their severity [40] or the location of the lesion [41]. The different predicted behaviours from our model can enable the grouping of patients according to the presumed neural mechanism of the lesion, based on their motor adaptation performance. For cerebellar ataxia patients, perhaps, the severity of ataxia, which is rated using the International Cooperative Ataxia Rating Scale [71], is actually driven by differences in the neural mechanism of the lesion. A hint for it is in the adaptation profile of the mild and severe patients in the study by Criscimagna-Hemminger et al [40], where the performance of severe patients was similar to lesion 3 in our model while the performance of the mild patients was similar to lesion 1. Deuschl et al [37] also had patients with mild cerebellar impairments, and their group results were also similar to lesion 1, while Maschke et al [38] had only patients with moderate to severe cerebellar ataxia and their group results were also similar to lesion 3, showing no aftereffects.

## Conclusion

We presented here a novel spiking neural circuit model of human motor adaptation experiments which spans multiple levels of biological organisation from neural activity in cerebellum to neuromechanics that reproduce and predict human experimental outcomes. We focused here specifically on the cerebellum and two learning mechanisms implemented as synaptic learning rules which allow us to model predictive learning and memory formation. In a force-field sensorimotor adaptation task, our model successfully reproduced learning curves, after-effects, and savings reported in human motor adaptation studies. The model successfully learns from motor perturbations (force field trials) and generate motor corrections while providing a bottom-up view for the neural basis of adaptation. Our results show that when we apply different lesions to the cerebellar circuits in our model it induced changes in the learning which are in-line with those reported in patients’ and animals lesions studies, thus show the potential to predict how specific types of cerebellar damage will affect patients’ learning capabilities. Models such as this presented here will become essential in closing the loop between circuit level mechanistic understanding, often obtained from small animal studies, to human brains and the full-body behaviour they control. This will be essential for linking the growing body of work in real-world neuroscience of studying human behaviour in real-live in real-live tasks [25, 72–75], to circuit level mechanisms – ultimately because the latter are under genetic and developmental control, but only the former determine the evolutionary fitness [76].

## Notes

https://doi.org/10.6084/m9.figshare.14837742.v1

### Competing Interest Statement

The authors have declared no competing interest.

