## Appendix for "Reproducing Human Motor Adaptation in Spiking Neural Simulation and known Synaptic Learning Rules"

#### Simulation of Force-field Reaching Task

We apply a group of parameters to describe the force-field task on a 2-dimensional space. To avoid unknown muscle kinematics and dynamics, we describe movement commands, perturbations and compensations all the form of hand accelerations.

- Actual body state $S = [H, V]$ with hand position $H = [h_x, h_y]^\top$ and hand velocity $V = [v_x, v_y]^\top$.
- Planned movement command $U^* = [u_x^*, u_y^*]^\top$, describing the acceleration of the hand caused by internal (muscular) force. It is pre-determined to follow a real-world human forward reaching movement (data from recorded non-perturbed movement in human experiments (**?**)).
- Perturbation $P = [p_x, p_y]^\top$, describing the acceleration of the hand caused by external force. All perturbations we applied in our force-field simulation are lateral perturbations along the x-axis, with amplitude proportional to the y-axis velocity:
$$P = [p_x, p_y]^\top = [bv_y, 0]^\top, \tag{1}$$
where $b$ is a perturbations factor setted for each trial ($b > 0/b < 0$ represents right/ left side perturbations).
- Compensation $C = [c_x, c_y]^\top$, generated by the system to cancel the influence of the perturbation.

Equations below describe the updating procedure of the hand state, driven by the internal motor command $U^*$, the external perturbation $P$ and the compensation $C$:

$$\begin{aligned} U(t) &= U^*(t) + P(t) + C(t), \\ V(t + \Delta t) &= V(t) + U(t)\Delta t, \\ H(t + \Delta t) &= H(t) + V(t)\Delta t + \frac{1}{2}U(t)\Delta t^2, \end{aligned} \tag{2}$$

where the executed motor command $U$ represents the overall acceleration of the hand caused by internal and external forces.

### System Design of Cerebellum-based Spiking Neural Network

The cerebellum-based spiking neural network involves a variety of neural populations, which is introduced in Fig. ?? with corresponding population size and neural model.

Input neurons in the cerebellum-based spiking neural network encode two signals: planned motor command and ($U^*(t) = [u_x^*(t), u_y^*(t)]$) and the planned movement ($S^*(t) = [p_x^*(t), p_y^*(t), v_x^*(t), v_y^*(t))$. They are Poisson neurons and encode information via rate coding. We separate input neurons into two groups for representing partial or full combination of input signals. The first group only encodes the planned motor command ($U^*(t)$) via population coding and project to the granule cell layer as motor efference. Each neuron has a nonlinear receptive field in the space of $U^*(t)$. The tuning function of i-th neuron with receptive centre $Z_i$ is given by the equation below.

$$h(U^*) = e^{\frac{(U^*-Z_i)^\top (U^*-Z_i)}{\sigma^2}} \tag{3}$$

where $\sigma$ is a constant controlling the expansion of the receptive field.

The second group of input neurons represents the complete upstream information ($[U^*(t), S^*(t)]$), which is a combination of 6 dimensions and not convenient to use the coding method in Eq. 3. It projects to the motor compensation neurons in the cerebellar nuclei for memory formation. We implement locationally-constrained (sparse) coding (LLC) to express the information in a relatively smaller size of neurons. LLC is an improvement of sparse coding (?) following the criteria:

$$\min_C \sum_{i=1}^{N} \|x_i - Bc_i\|^2 + \lambda \|d_i \odot c_i\|^2, s.t. 1^T c_i = 1, \forall i \tag{4}$$

where $\odot$ means the multiplication for each corresponding pair of elements in matrix and $d_i$ is the locality adapter based on distance. Compared with the standard sparse coding, the add-on item of the coding criteria ($\lambda \|d_i \odot c_i\|^2$) guarantees locality in representation and smooth neural activity changes when the upstream information varies gradually.

Action potentials of Poisson input neurons are delivered to the cerebellar cortex and the cerebellar nuclei via mossy fibres. The cerebellar cortex involves a liquid state machine with a recurrent neural network (the granule cell layer) and read-out neurons (the Purkinje cell layer). The recurrent synapse in the granule cell layer leads to complex neural dynamics depending on a series of current and previous inputs. It provides firing patterns containing spatial-temporal information, which is read by the Purkinje cell layer via a feedforward projection (synapses). We apply two groups of Purkinje cells to decode multiple information: the position $h_x$ and the velocity $h_y$. The neural coding scheme for Purkinje cells are population coding with predetermined receptive fields. Each group self-connected to perform competition, making sure only a small group of neurons with similar code (or receptive fields) are activated. The error-based teaching signal for predictive learning arises from the inferior olive via climbing fibres. The neural circuits around the inferior olive is still unclear. Thus, we define this teaching signal as the difference between the Purkinje layer estimation and the sensory feedback, namely the predictive error if the sensory system is accurate and unbiased.

Corresponding to the Purkinje cell layer, the population of movement error neurons are divided into two sub-groups, representing the lateral deviation and the differentiation of deviation. The motor compensation neural population is the only output of the cerebellum-based spiking neural, sending motor compensation commands to change body movement under perturbations. It receives inputs from movement error neurons as well as the experience-dependant memory. This error-compensation subsystem performs proportional-derivative (PD) control, providing instantaneous response to the control error and the derivative of change of the control error.

Synaptic weights in the spiking neural network are pre-defined and fixed during the entire force-field simulation except for the parallel fiber-PC synapses and the mossy fiber-ME synapses. These two synapses are initialised with random weights and pre-trained in 30 baseline force-field trials to allow the formation of appropriate synaptic connections via synaptic learning rules.
